# Dynamic Lateralization in Contralateral-Projecting Corticospinal Neurons During Motor Learning

**DOI:** 10.1101/2024.02.08.579456

**Authors:** Jiawei Han, Ruixue Wang, Minmin Wang, Zhihua Yu, Liang Zhu, Jianmin Zhang, Junming Zhu, Shaomin Zhang, Wang Xi, Hemmings Wu

## Abstract

Understanding the adaptability of the motor cortex in response to bilateral motor tasks is crucial for advancing our knowledge of neural plasticity and motor learning. Here we aim to investigate the dynamic lateralization of contralateral-projecting corticospinal neurons (cpCSNs) during such tasks. Utilizing in vivo two-photon calcium imaging, we observe cpCSNs in mice performing a bilateral lever-press task. Our findings reveal a heterogeneous population dynamics in cpCSNs: a marked decrease in activity during ipsilateral motor learning, in contrast to maintained activity during contralateral motor learning. Notably, individual cpCSNs show dynamic shifts in their engagement with ipsilateral and contralateral movements, displaying an evolving pattern of activation over successive days. This suggests a flexible and learning-related reconfiguration in the cpCSN network, parallel to motor learning stages. Our findings suggest a complex reorganization in cpCSN activities, underscoring the dynamic nature of cortical lateralization in motor learning and offering insights for neuromotor rehabilitation.

## Introduction

The motor cortex (MC) is considered to play an important role in motor skill learning^1^. Motor skill learning is compromised when the integrity of MC is impaired^2–6^. Plasticity at multiple levels of the MC is implicated during motor learning, including changes in dendritic spines and neuronal activity in different layers, inputs from subcortical structures, and stimulation threshold and cortical mapping to evoke movement^7–13^.

Two types of learning-related plasticity are reported in the MC. On the one hand, the activity of the MC neurons is closely correlated with movement during learning^14,15^. On the other hand, the relationship between the MC neuronal activity and movement is highly dynamic^10,15–17^. Furthermore, it has been shown that the role of the motor cortex in the execution of well-learned motor skills may become less critical^2,18^. This duality in plasticity represents a balance between stable neural patterns, which underpin the precise execution of well-learned motor skills, and dynamic neural adaptations, which allow for learning of new movements and adjustment of existing ones, illustrating the MC’s pivotal role in both the acquisition and refinement of motor skills. This balance ensures the motor system’s adaptability and efficiency, facilitating both the consolidation of motor memory and the flexibility required for continuous learning. Such a duality has also been reported in the plasticity of the corticospinal neuron (CSN), the main output of the MC in layer 5 during motor learning^8,14,16,17,19,20^. It has been suggested that the activity of CSN is highly consistent with movement in the limbs. This conjecture has been confirmed in recent studies that the consistency between CSN activity and movement is better than that of layer 2/3 neurons in the MC. However, CSN also shows dynamic plasticity during motor learning, and the relationship between an individual CSN and movement fluctuate with learning^9,17^. Approximately 90% of the CSNs terminate in the contralateral spinal cord to form a single synaptic connection^17,21^, while sending axon branches to the ipsilateral subcortical structures, pons and bilateral spinal cord at upper levels^21,22^. These contralateral-projecting CSNs (cpCSNs) are the major output of regular spiking neurons in layer 5, and regular spiking neurons encode bilateral movement information. Whether and how cpCSNs modulate bilateral movement remains unclear.

Here we aim to investigate the dynamics of cpCSNs representation of bilateral movement during motor learning. First, we investigate whether cpCSNs carry ipsilateral movement information, and second, if such is the case, how cpCSNs distribute workload collectively and maximize efficiency during motor learning.

## Results

### Two-photon calcium imaging of cpCSNs during a bilateral motor task

To investigate the dynamics of contralateral-projecting CSNs (cpCSNs) in ipsilateral and contralateral movement during learning, we used the Cre-FLEX system to selectively image cpCSNs under a two-photon microscope. This was achieved by a combination of adeno-associated virus encoding the calcium indicator GCaMP6f (AAV2/9-EF1α-DIO-GCaMP6f) injection in the left motor cortex and AAV encoding Cre recombinase (AAV2/9-CaMKII-Cre) injection into the C6 to C8 segments of the right spinal cord (Figure 1A; STAR Methods). Fluorescence of cpCSNs was observed 2 weeks after injection, however, long-term imaging of somas in layer 5 was not possible due to their depth. Instead, we imaged apical dendritic trunks in layer 2/3, as their calcium events showed a high correlation with those of the somas, serving as a reliable proxy. To stably record the activity of dendritic trunks, we used vessels in the field of view (FOV) for image alignment (Figure S1).

**Figure 1.**
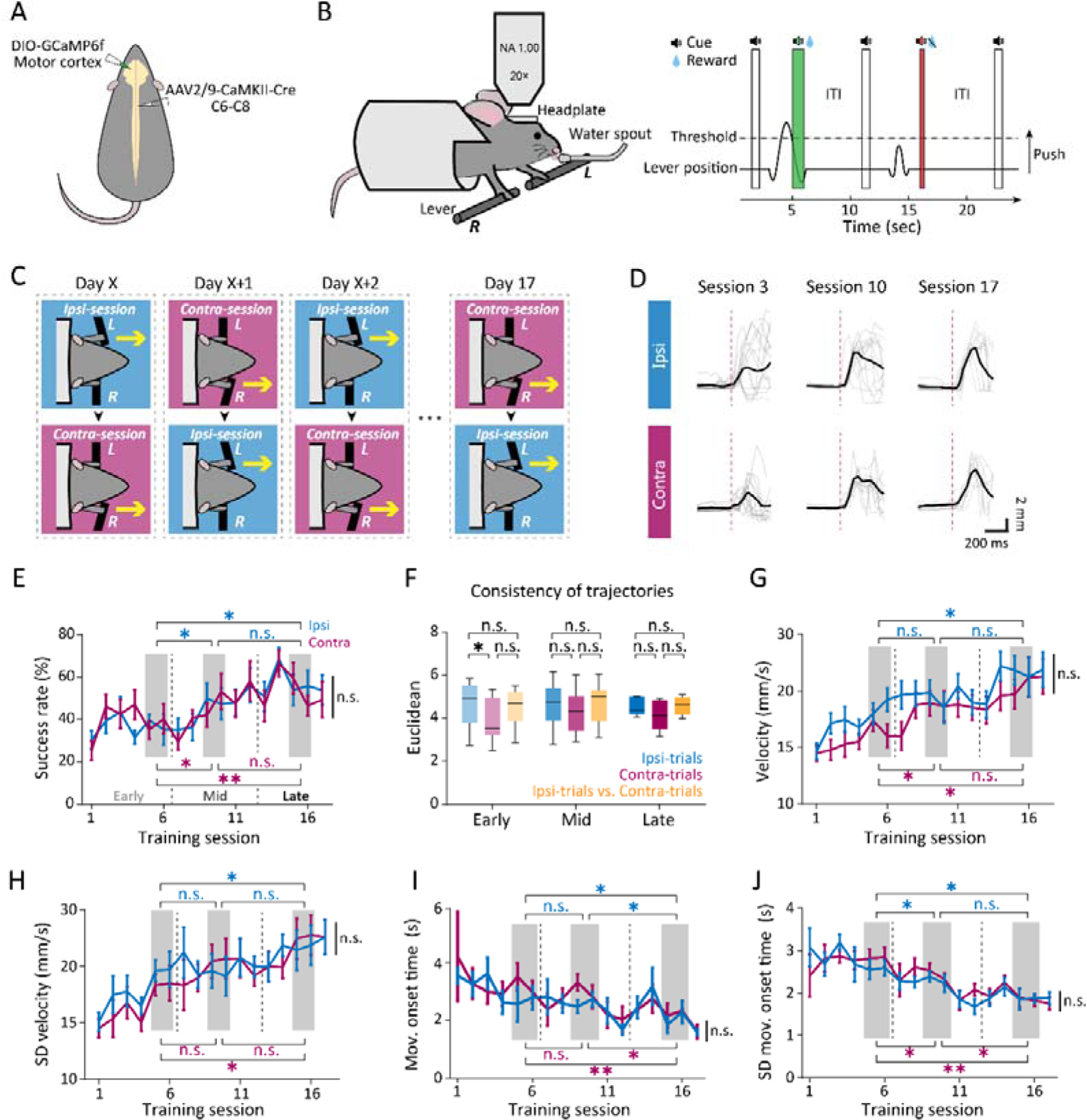
Virus injection and the bilateral lever-press task. (A) Schematic of injections to selectively express GCaMP6f in corticospinal neurons (cpCSNs). (B) Behavioral setup of the ipsilateral and contralateral lever-press task. Left: the mouse was trained to push the left (or right) lever with the corresponding forepaw for a water reward. Right: passing the lever position threshold after a start cue (black horn and white bar) resulted in a water reward (blue droplet) and a success cue (green horn and bar), or in white noise (red horn and bar) without a reward. ITI: inter-trial interval. (C) The task consisted of ipsilateral or contralateral sessions each day, alternating daily. (D) Lever trajectories of sessions 3, 10, and 17 in an example mouse. Top: ipsilateral trials. Bottom: contralateral trials. Gray lines: single movements from randomly selected 15 rewarded trials; black lines: average of these trials; red dashed line: movement onset. (E) Success rate comparison between ipsilateral and contralateral trials showed no significant difference (p > 0.05, two-way repeated-measures ANOVA, n = 10 mice from sessions 1 to 17, mean ± SEM). Success rates improved over the course of learning (comparison with average values at early (sessions 5, 6), mid (sessions 9, 10), and late (sessions 15, 16) learning stages; n = 10 mice; *p < 0.05, **p < 0.01, n.s., not significant, Mann-Whitney U test). (F) Euclidean distances within ipsilateral trials, contralateral trials, and between trial types at different learning stages (significant difference noted at the early stage between ipsilateral and contralateral trials, p < 0.05; n.s., not significant, two-sample t-test, mean ± SEM). (G-J) Analyses of mean velocity, variability in velocity, movement onset time, and variability in movement onset time revealed improvements over learning sessions, and no significant differences between ipsilateral and contralateral trials (n.s., not significant, *p < 0.05, **p < 0.01, p values from two-way repeated-measures ANOVA and Mann-Whitney U tests as appropriate, mean ± SEM).

Over a period of 3 weeks, mice (n = 9) were trained daily in both an ipsilateral session and a contralateral session on a bilateral lever-press task (Figure 1B). This task was adapted from traditional lever-press and Right-Left Pedal tasks^23,24^. The bilateral lever-press task consisted of two parts to dissociate the movement of ipsilateral and contralateral limbs so that neuron activity could be recorded respectively. In the ipsilateral session, water-restricted mice learned to use their left (ipsilateral) forelimb to press the left lever for a water reward, and conversely, used their right (contralateral) forelimb in the contralateral session. The duration of each session was 10 minutes, and the order of ipsilateral and contralateral session was reversed every day (Figure 1C). Lever trajectories on both sides became more consistent with training over time (Figure 1D). The lever press performance—measured by success rates, consistencies, velocities, and response times—improved significantly during motor learning in both ipsilateral and contralateral movement, and showed no significant difference between the two sides (paired t-test, p > 0.05; Figures 1E-J). These results suggested that mice were able to learn bilateral lever-press task at similar level.

### Neural representation of cpCSNs during bilateral movements

We then investigated neuronal activity of cpCSNs during ipsilateral and contralateral movements. Similar to previous findings in motor cortical neuron, some cpCSNs were active during movements, while others remained inactive (Figure 2A)^9,17,18,24,25^. We categorized cpCSNs into task-related (active) neurons and silent neurons based on their calcium events during movements (STAR Methods). Additionally, active cpCSNs exhibited heterogeneity during bilateral movements. Neurons active solely during ipsilateral movements were classified as ipsilateral-preferring (ipsi-preferring) neurons. In contrast, neurons active only during contralateral movements were classified as contralateral-preferring (contra-preferring) neurons. Neurons active and inactive during both ipsilateral and contralateral movements were classified as bilateral-preferring (bi-preferring) neurons and silent neurons, respectively (Figures 2B and 2C and Figure S2; STAR Methods). These findings suggested that cpCSNs were involved in the control of both ipsilateral and contralateral movements.

**Figure 2.**
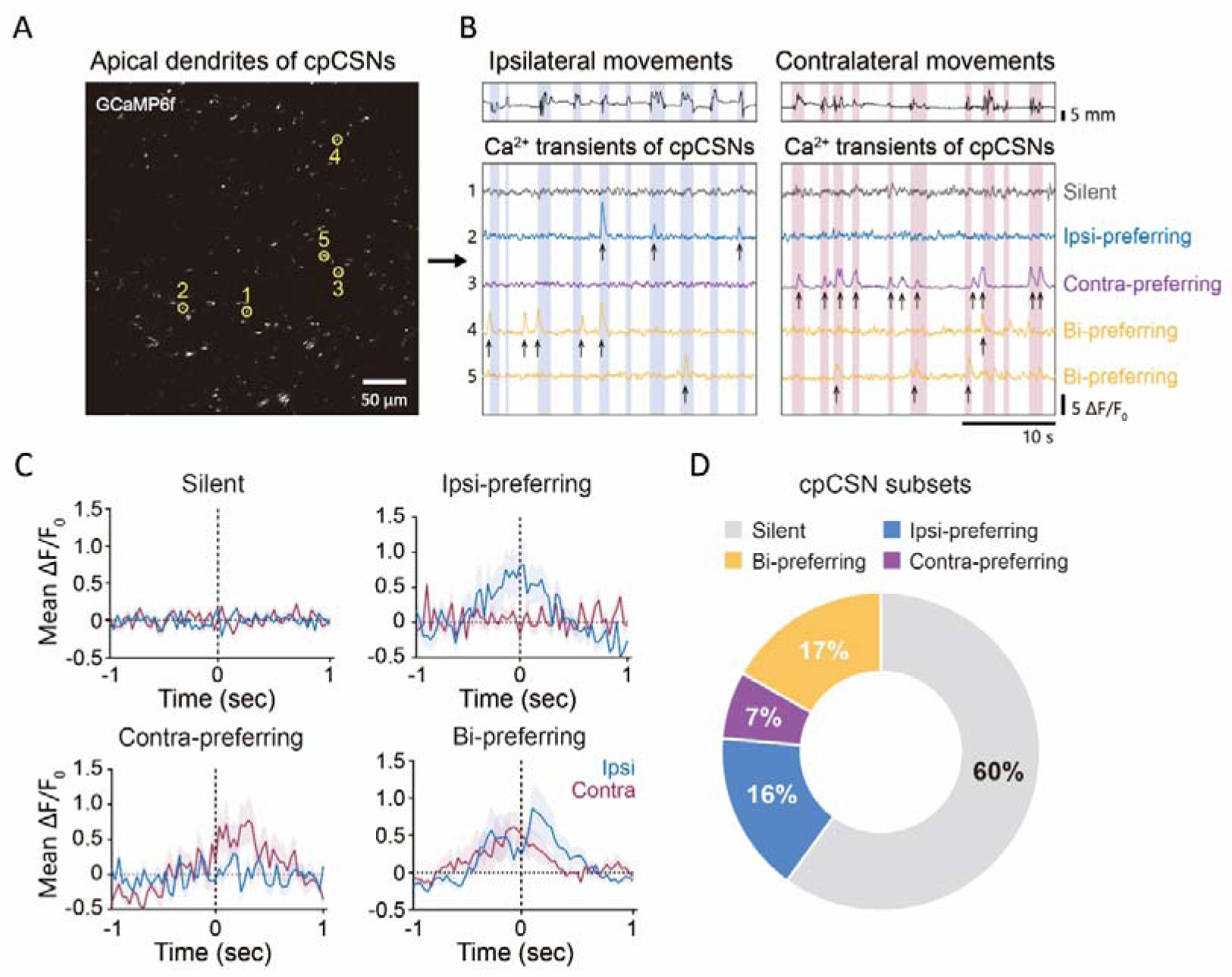
Heterogeneity of cpCSNs in Ipsilateral and Contralateral Movements. (A) Two-photon (2p) image showing GCaMP6f expression in apical dendrites of contralateral-projecting corticospinal neurons (cpCSNs) with example regions of interest (ROIs) highlighted in yellow. (B) Activity of the example ROIs from (A) during movements in a representative mouse. Top: lever position for ipsilateral and contralateral movements, with green highlighted regions indicating detected movements. Bottom: calcium transients of silent, ipsi-preferring, contra-preferring, and bi-preferring cpCSNs, with arrows marking detected calcium events. (C) Mean ΔF/F_0_ of example cpCSNs subsets across 30 randomly selected ipsilateral (ipsi) trials and 30 contralateral (contra) trials (mean ± SEM). (D) Proportional distribution of early-stage cpCSN subsets.

### The dynamics of laterality in cpCSN ensembles during motor learning

To elucidate the dynamics of cpCSNs during motor learning, we quantified the proportions of ipsi-, contra-, bi-preferring, and silent neurons at different stages of learning. We found no significant changes in the proportions of silent neurons (early: 60.03 ± 4.51%, mid: 54.72 ± 5.7%, late: 47.05 ± 4.63%; mean ± SEM), and bi-preferring neurons (early: 16.78 ± 3.37%, mid: 21.4 ± 4.96%, late: 22.5 ± 5.33%; mean ± SEM; Figure 3A) across stages. However, the proportion of ipsi-preferring neurons was significantly higher in the early and late compared to the middle stage (early: 16.31 ± 1.53%, mid: 8.81 ± 1.23%, late: 17.83 ± 2.79%; mean ± SEM; Figure 3A), while the proportion of contra-preferring neurons was significantly lower in the early stage relative to the middle and late stages (early: 6.88 ± 1.03%, mid: 15.04 ± 2.33%, late: 12.57 ± 2.06%; mean ± SEM; Figure 3A). These alterations in the proportions of ipsi- and contra-preferring neurons suggested laterality-related plasticity within cpCSN ensembles during bilateral motor learning.

**Figure 3.**
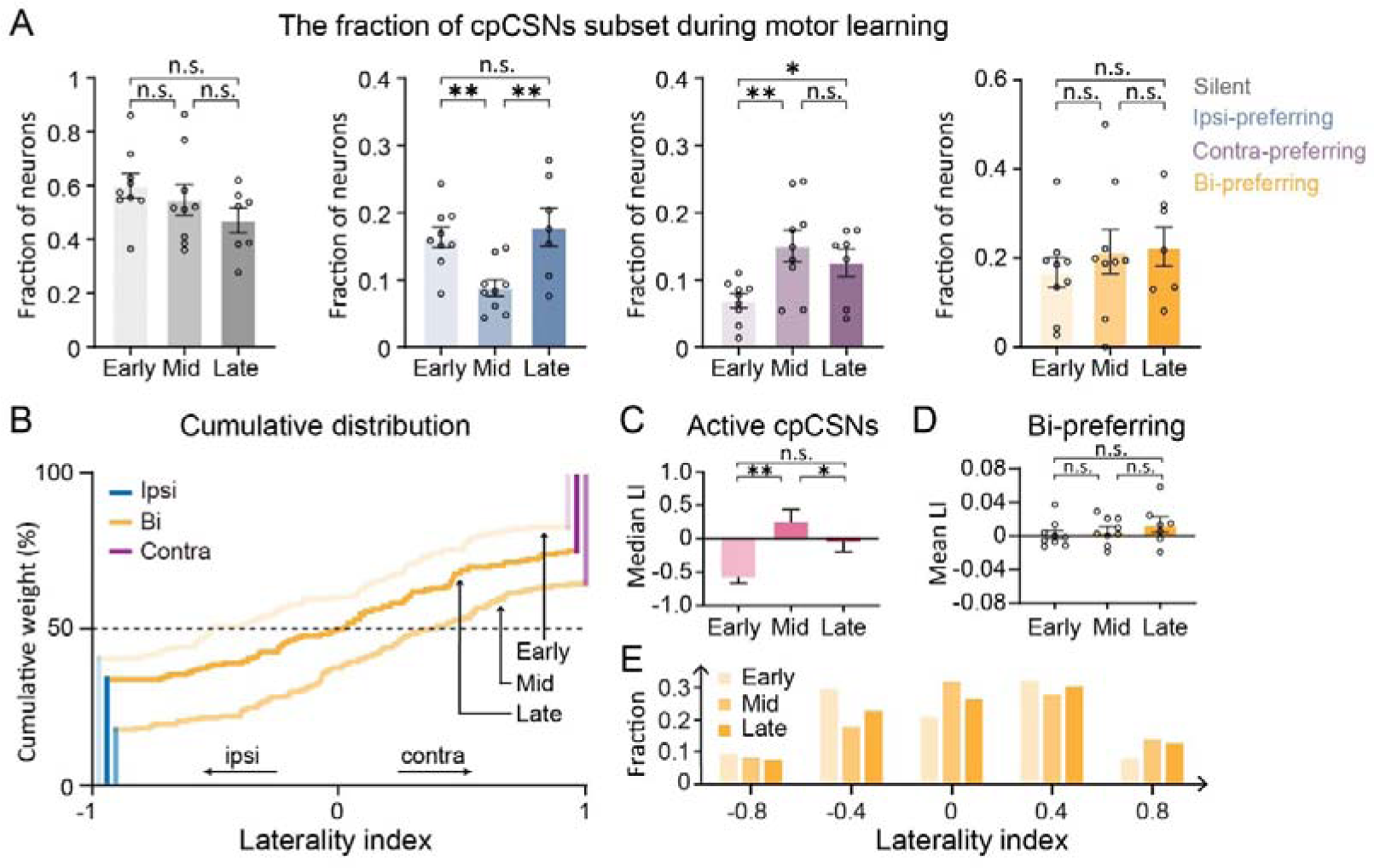
Dynamics of cpCSN Laterality During Motor Learning. (A) Proportions of classified cpCSNs at different learning stages (n = 9 mice, 548 neurons in total, n.s., not significant, *p < 0.05, **p < 0.01, paired sample t-test, mean ± SEM). (B) Cumulative distribution of the laterality indices (LI) for all active cpCSNs at the early, middle, and late stages of learning. (C) Median LI of all movement-related cpCSNs across different stages (n = 548 neurons, n.s., not significant, *p < 0.05, **p < 0.01, independent-sample t-test, mean ± SEM). (D) Mean LI of bi-preferring neurons at different stages (n.s., not significant, paired sample t-test, mean ± SEM). (E) Distribution of bi-preferring neuron laterality indices, ranging from −1 to 1, divided into five equal groups according to the laterality index, at different stages.

The cumulative distribution of the laterality index among the cpCSN ensemble revealed an ipsilateral preference at the early stage, a contralateral preference at the middle stage, and no discernible preference at the late stage (Figure 3B). The mean laterality index at the level of individual mice followed a similar pattern (laterality index: early: −0.57 ± 0.1, mid: 0.24 ± 0.19, late: −0.04 ± 0.15; mean ± SEM; Figure 3C). To determine whether these changes in ensemble laterality were attributable to shifts in ipsi- and contra-preferring neurons or bi-preferring neurons, we analyzed the mean laterality index of bi-preferring neurons at different stages. No significant changes were found between stages (laterality index: early: 0.002 ± 0.005, mid: 0.006 ± 0.005, late: 0.013 ± 0.009; mean ± SEM; Figures 3D and 3E), indicating that the dynamics of ipsi- and contra-preferring neurons were the primary contributors to the laterality dynamics in cpCSNs.

### Laterality-dependent plasticity of cpCSNs during motor learning

Based on these findings, it appeared to us that the changes in the percentages of ipsilateral- and contralateral-preferring cpCSNs were complementary to each other (i.e. there was an increase in contralateral-preferring and a decrease in ipsilateral-preferring neurons from the early to the middle stage, and vice versa from the middle to the late stage). To further confirm this hypothesis, we investigated the activities of individual neurons during ipsilateral and contralateral motor learning tasks. The laterality classifications of individual cpCSNs from the early, middle, and late stages were shown in Figures 4A-D. We found that only 26.8% of the ipsilateral-preferring neurons in early stage changed their preference to contralateral in middle stage (Figure 4E). Similarly, only 16.6% of the contralateral-preferring neurons in middle stage changed their preference to ipsilateral in late stage (Figure 4F). These findings indicated that the changes we observed at the population level may not accurately reflect individual neuronal dynamics. At the level of individual neuron, only nine of the 67 bi-preferring cpCSNs (13.4%) and 39 of the 212 silent cpCSNs (18.4%) remained unchanged throughout the motor learning task. The majority of the cpCSNs, including 86.6% of the bi-preferring and 81.6% of the silent cpCSNs, as well as all of the ipsilateral- and contralateral-preferring cpCSNs, changed their laterality preference (Figure 4G). The laterality dynamic index suggested that silent cpCSNs were the most stable group of neurons in terms of laterality preference, while ipsilateral-preferring cpCSNs showed a significantly higher dynamic index (Figure 4H). The cpCSNs were categorized based on the number of changes in laterality preference, as summarized in Figure 4I, suggesting that more than 85% of cpCSNs changed preference between every learning session.

**Figure 4.**
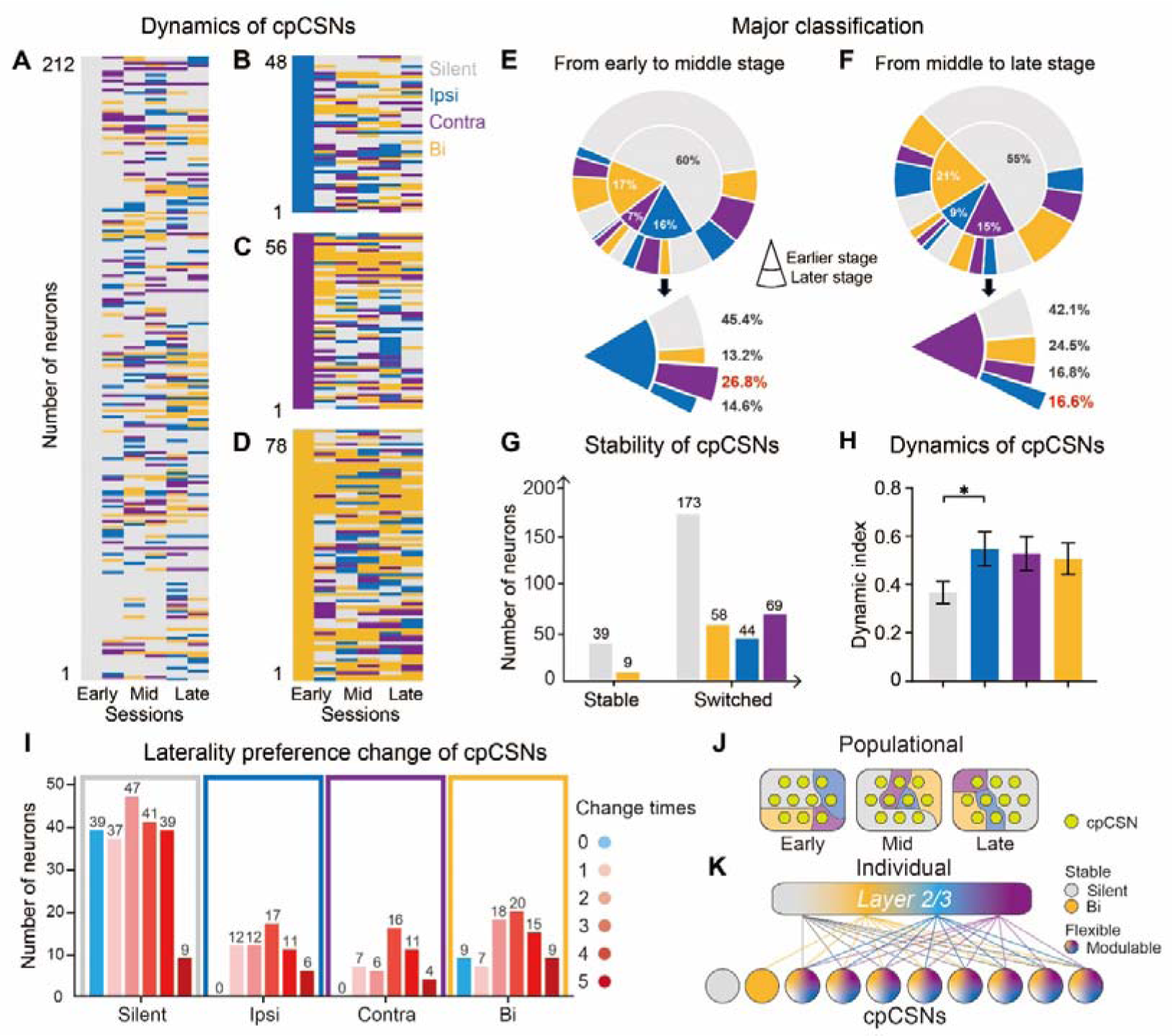
Versatility of laterality-dependent plasticity in cpCSNs during learning. (A-D) Laterality classifications of individual cpCSNs across learning sessions: silent (A), ipsilateral-preferring (B), contralateral-preferring (C) and bi-preferring (D) neurons in the first session. (E-F) Major classification (the classification with the largest number of days at one stage) of cpCSNs from early to middle stage, and from middle to late stage. (G) Number of cpCSNs that maintained or changed their classification in laterality preference during learning. (H) Dynamic index of cpCSN subsets during learning (*p < 0.05, two independent-samples t-test, mean ± SEM). (I) Number of cpCSN subsets and the frequency of changes in their laterality preference during learning. (J-K) Schematic representations of the dynamic lateralization of cpCSNs at population (J) and individual neuron (K) levels during motor learning. At the population level, the proportions of cpCSN subsets, based on their laterality preferences, dynamically change across different stages of learning. At the individual neuron level, apart from a small subset of silent and bi-preferring cpCSNs, the majority exhibit high laterality modulability during motor learning, indicating day-to-day changes.

## Discussion

### Corticospinal Neurons in Motor Learning

Corticospinal neurons, especially contralateral-projecting corticospinal neurons (cpCSNs), play a crucial role in the diverse modulation observed during motor learning, as evidenced by previous studies^17,20,26–29^. Through a bilateral lever-press task, we tracked the activities of cpCSNs’ apical dendrites in both ipsilateral and contralateral movements throughout motor learning. Our results uncover dynamic shifts in laterality preferences and proportions of cpCSN subsets over the learning trajectory A majority of individual cpCSNs displayed malleable laterality preferences, suggesting intricate interactions between cpCSN activities and the direction of movement.

### Dynamics in cpCSN ensemble

The neuronal activity of cpCSNs is selectively active during ipsilateral or contralateral movements (Figure 2), which highlights the heterogeneity in their laterality preferences, and aligns with previous research^17,18,23,24^. In the motor cortex, pyramidal tract neurons exhibit task-related activity with distinct laterality preferences in newly learned motor tasks^24^. Furthermore, individual cpCSNs displayed diverse activity patterns, switching between movement-active and quiescence-active during the learning process^17^. The capacity of cpCSNs to represent bilateral motor information concurrently parallels previous discoveries that neural signals in one hemisphere can decode movements of both forelimbs^30^. The functional diversity of cpCSNs may be linked to their axonal lateral branches, which are responsive to different neurotransmitters and undergo distinct modulation during learning periods^31,32^.

We categorized cpCSNs into four subsets based on laterality preferences. The subset proportions shifted during motor learning (Figure 3), consistent with changes in movement-active and movement-related neuron proportions previously reported^17,18^.

Intriguingly, the later learning stages saw a surge in ipsi-preferring neurons, diverging from studies reporting a dominance of contra-preferring pyramidal tract neurons^24,33^. This divergence might stem from our study’s focus on cpCSNs, rather than previously studied contralateral-projecting corticopontine neurons. Additionally, our larger neuron sample size could contribute to this discrepancy. Task-specific neuronal adaptations to varied motor tasks further suggest that cpCSNs serve multifaceted roles during different learning stages^26,34,35^.

### Motor Cortex Laterality Preference: A Dynamic Output

The dynamic nature of cpCSNs’ laterality preference throughout the learning stages is a key aspect of our findings (Figures 4J and 4K). Individual cpCSNs displayed significant flexibility, with transitions between movement-active and quiescence-active states, indicative of a bidirectionally evolving pattern of activity^17,18^. The motor cortex’s task-specific patterns surpass the activity differences due to muscle contraction alone^19,36,37^, reflecting neural ensembles’ capacity to buffer against input noise^17,38–40^. Additionally, our results show that the activity patterns in cpCSNs are adaptable during both ipsilateral and contralateral movements in the learning process.

Remarkably, the increase in the proportion of ipsi-preferring neurons from the middle to the late stage of learning did not result from a corresponding decrease in contra-preferring neurons. Instead, this increase comprised changes across all four cpCSN subsets. The primary axons of cpCSNs, which terminate in the contralateral spinal cord, suggest a dominant role in controlling movements of contralateral limbs^41,42^. However, cpCSNs also develop axon collaterals that project to subcortical structures on the ipsilateral side, particularly the dorsal striatum^26^. It has been shown that spiny projection neurons (SPNs) in the dorsal lateral striatum, which receive input from the motor cortex, exhibit synaptic plasticity to both ipsilateral and contralateral movements during learning^8^. This indicates that cpCSNs, as precursors to SPNs, may exhibit plasticity in response to bilateral movements during learning phases. Furthermore, the motor task’s stereotyped pattern leads to motor skill automatization by subcortical structures, rather than the motor cortex itself^2,18,25^, reducing cortical interference during movement execution^43^.

In conclusion, our study highlights the modulability of cpCSNs in response to laterality preference during bilateral motor learning. These findings offer insights into the potential roles of cpCSNs in motor rehabilitation, demonstrating their capacity to adapt dynamically to varying learning stages and tasks. The versatility and adaptability of cpCSNs underscore their importance in the neural mechanisms underlying motor learning, and provide a foundation for future research into their role in motor skill acquisition and rehabilitation.

## Methods

### Specimens and ethics

All animal studies and experimental procedures were approved by the Animal Care and Use Committee of the animal facility at Zhejiang University and in accordance with the Institutes of Health Guide for the Care and Use of Laboratory Animals. A total of 10 adult male mice (C57BL/6J, 8 weeks or older) were used in the experiments. Mice were grouped and housed in a 12:12 hours reverse light-dark cycle, and all experimental activities were conducted during the dark phase to maintain consistency.

### Surgeries and virus injections

Surgical procedures were performed under isoflurane anesthesia (4% for induction and 1.5-2% during surgery) and injected with dexamethasone (2 mg/kg), cefuroxime (5 mg/kg) and buprenorphine (0.1 mg/kg) subcutaneously at the beginning of the surgery to prevent infection, inflammation, and pain. Surgeries consisted of two successive steps, the first being a spinal cord injection, and the second being a cortical injection and cranial window implantation, as described by Peter et al.^17^.

For the first step, the back was shaved and cleansed with iodine, alcohol, and saline. A midline incision was then made in the skin, reaching deep into the muscle layer. Muscle was removed and fatty tissue was isolated to fully expose the thoracic vertebrae. Because the spinous process of T2 was prominent, the muscles and ligaments attached to the spine of the C5-T2 segment were located and cleared. The T2 spinous process was clamped and fixed, and the soft tissue on the surface of the C6-C8 segments of the spinal cord was removed. AAV2/9-CaMKII-Cre (BrainVTA Technology, China) inside a glass electrode was injected into three sites on the right side of the spinal cord (200 nL at each site) and placed 0.4 mm from the midline, 0.7 mm from the surface and separated by 0.6 mm rostrocaudally. After the injection, the fixed T2 spinous process was released and the muscle, soft tissue, and skin were sutured sequentially. Subsequently, for cortical injections, we prepared the skull surface and opened a cranial window over the left motor cortex, injecting AAV2/9-EF1α-DIO-GCaMP6f (BrainVTA Technology, China) at five sites in a plus (+) shape in the left motor cortex, each injection was 50 nL and placed 0.7 mm from the surface, separated by 0.5 mm, centered at the left caudal area (0.3 mm anterior and 1.5mm lateral to Bregma). A 5 mm-diameter glass coverslip was placed over the skull window and secured with medical glue and dental cement. A custom-made metal headplate was then glued to the skull surface and fixed with dental cement. Post-operative care ensured animals displayed no motor deficits, with most resuming wheel running within two days.

### Behavior

After recovering from surgery, mice were restricted to a maximum of 1-2 mL water per day for 2 weeks before the training. Mice were then trained daily in a lever-press task during two-photon imaging for 1 ipsilateral session and 1 contralateral session, lasting 10 mins per session. During lever-press task, mice rested their body and hindlimbs in a tube and placed their right forelimb on the fixed right lever and their left forelimb on a movable lever in the ipsilateral session, and vice versa in the contralateral session. The lever consisted of a handle glued to a custom angle encoder. Angle data from the angle encoder was continuously monitored using a customized decoder and software (The QT Company). Presses were defined as the lever being pushed beyond 6 mm and maintained for more than 120 ms before returning to the resting position. The task consisted of a variable intertrial interval, followed by a 5-10 s cue period during which the lever pressed triggered water reward was delivered. Cue periods paired with a 1-second start cue, rewards paired with a 600-ms high-pitched melodious tone, and a failure to press the lever within the cue period paired with a 200-ms white noise, with an extra 2 s of intertrial interval. The cue period was reduced during the first three sessions from 15 s to 10 s and then to 5 s, and the intertrial interval was increased during the first 5 sessions from 2-3 s to 5-7 s and then to 8-10 s to encourage discrete movements. The duration time was increased from 60 ms to 120 ms and the lever-press threshold was increased from 3 mm to 6 mm during the first four sessions. Each session consisted of approximately 80 trials. Sessions were binned into 3 stages in analyses (early: 1-6; middle: 7-12; late: 13 and after). At the early stage of learning, various behavioral parameters were constantly adjusted to enhance learning until the 5th session, after which the behavioral parameters would remain fixed. The average success rate exceeded 50% on session 12, and as a result, training after session 12 was defined as the late stage (Figure S3).

For all mice, if an ipsilateral session was performed first on one day and a contralateral session was performed after a rest of 5-10 mins, the contralateral session was performed first and then the ipsilateral session was performed on the next day, to prevent the corresponding laterality of mice due to task stereotype.

### Lever-press movement analysis

The angle data were down-sampled from 200 Hz to 30 Hz to remove baseline noise. The movement onset time for each lever press was determined by finding when the lever moved away from the baseline position before movements. The movement epoch was defined as −1 s to 1 s around the movement onset. The largest lever angle and the time duration for the lever pushed beyond 5 mm were used as the motion features to further filter out the middle 80% of movements within each session. All filtered lever presses were aligned by the movement onset time for subsequent analyses.

We used Euclidean distance (d) to describe the consistency between different lever trajectories:

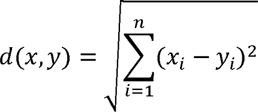

Where trajectory A (x_1_, x_2_,…, x_n_) and trajectory B (y_1_, y_2_,…, y_n_) are the normalized lever position during the movement-related period from two different trials. Euclidean distance between each pair of trajectories from different trials within session or across sessions comprising the mean distance of the sessions were calculated.

### Two-photon imaging

Before imaging, one dose of 50 μL TRITC-dextran solution (ALADDIN, China) was retro-orbitally injected. Two-photon imaging was conducted through a 20× 1.0 NA (Olympus) mounted on a customized two-photon microscope (2P plus, Bruker Corporation) with a tunable femtosecond laser (Ti: Sapphire laser, Chameleon Ultra II, Coherent Inc.), whose power was controlled by Pockels Cell (EO-PC, Thorlabs Corporation). Dual-channel fluorescent signals (green: 500-550 nm filter, red: 570-620 nm filter) were collected in photomultiplier tubes (PMTs). The wavelength of the imaging laser was first kept at 860 nm with low power (10-50 mW, under objective), so that imaging drifts could be manually monitored and corrected via identification of local vasculature (red channel) and apical dendrites (green channel). Then images were acquired at a rate of ∼30 Hz, covering 406.36×406.36 μm with 512×512 square pixels, using a 920 nm laser with low power (10-50 mW, under objective) for 10 mins per session to record the activity of apical dendrites (green channel).

For 3D structural imaging, the field of view was the same size at a rate of ∼0.5 Hz with 1024×1024 square pixels, in which each frame was repeated two times on average as structural results.

### Two-photon data analysis

The fluorescence data acquired by the two-photon microscope was first processed by Suite2p^44^, including motion correction, region of interest (ROI) generation, and fluorescence extraction. ROIs were then manually corrected by average image and fluorescent traces.

The average image of each session after motion correction was used to align the imaging data over multiple sessions. The average image of the first session in each mouse was picked as a reference, and the others were aligned to it via vasculature landmarks to perform normalized cross-correlation, and to find the coordinates of the peak. The alignment across sessions was based on the offset found by this correlation. In aligned maps, ROIs were evaluated by the proximity of centroids. If the distance between centroids of the candidate ROI was less than 8.5 pixels, the candidate ROI would be considered the same ROI. Only ROIs stably identified on given imaging sessions were included for all of the following analyses.

Fluorescent traces for each ROI were created by averaging enclosed pixels and subtracting background fluorescence.

The relative change in fluorescent traces for each ROI was using the following formula:

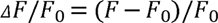

The F_0_ was estimated as the baseline of fluorescent traces from a ± 15 s sliding window. Then ΔF/F traces of each ROI were aligned by movement onset time, and then averaged across trials.

Ca events were calculated by ΔF/F_0_ traces for each ROI^17^. Two thresholds were defined to find active portions and baseline, the former was 3× standard deviation and the latter was 1× standard deviation. Active portions were identified by a 1-s LOESS-smoothed trace crossing the active threshold and extended backward to begin when the baseline threshold was last crossed. Periods with negative slopes during active portions were regarded as inactive. The remaining active portions were characterized as Ca events and set to the difference between the maximum and minimum values within each event, with baseline and inactive portions set to zero.

### Movement-related classification

cpCSNs were classified as movement-related (active) neurons or silent neurons each day. The neuron was classified as a movement-unrelated (silent) neuron on one day if the count of calcium events is less than 5 during movement epochs in both ipsilateral and contralateral sessions.

### Laterality index

To evaluate the laterality of each active neuron, we calculate calcium events per frame averaged over trials during movement, between ipsilateral and contralateral sessions. The laterality index (LI) was computed using the following formula:

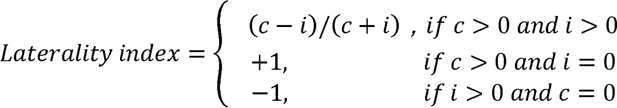

Where *c* and *i* are mean calcium events during contralateral and ipsilateral trials (−1 to 1 s relative to the onset of lever-press), respectively. If this index is 1, the neuron was classified as a contra-preferring neuron. If it is −1, the neuron was classified as an ipsi-preferring neuron. Otherwise, it was classified as a bi-preferring neuron.

### Dynamic index

To evaluate the dynamic strength in cpCSNs, we investigated their daily classifications. When the classification of cpCSNs shifts, we marked its change as 1, otherwise as 0. The dynamic index is the sum of the number of classification change divided by the number of sessions minus one.

### Statistical analysis

All statistical analyses were performed using IBM SPSS. The comparison of successful trials and successful rates between ipsilateral and contralateral movement were analyzed using two-way repeated-measures ANOVA. The laterality index of neurons, and the average activity of neurons at different stages during motor learning were analyzed using Mann-Whitney tests. The laterality index of neurons in different stages and the activity of neurons between ipsilateral and contralateral movement in the same stage were analyzed with paired sample t-tests. All comparisons using t-tests are two-sided. Differences were considered statistically significant when p < 0.05. Error bars indicate standard errors of the mean (SEM) unless noted otherwise.

## Notes

### Competing Interest Statement

The authors have declared no competing interest.

