## Supplemental Figure 1 for "Dynamic Lateralization in Contralateral-Projecting Corticospinal Neurons During Motor Learning"

**Figure S1. Imaging and alignment of contralateral-projecting corticospinal neurons (cpCSNs).**

**Figure S2. Examples of activities of contralateral-projecting corticospinal neurons (cpCSNs) during movements.**

**Figure S3. Behavior task setup in ipsilateral and contralateral sessions during motor learning.**

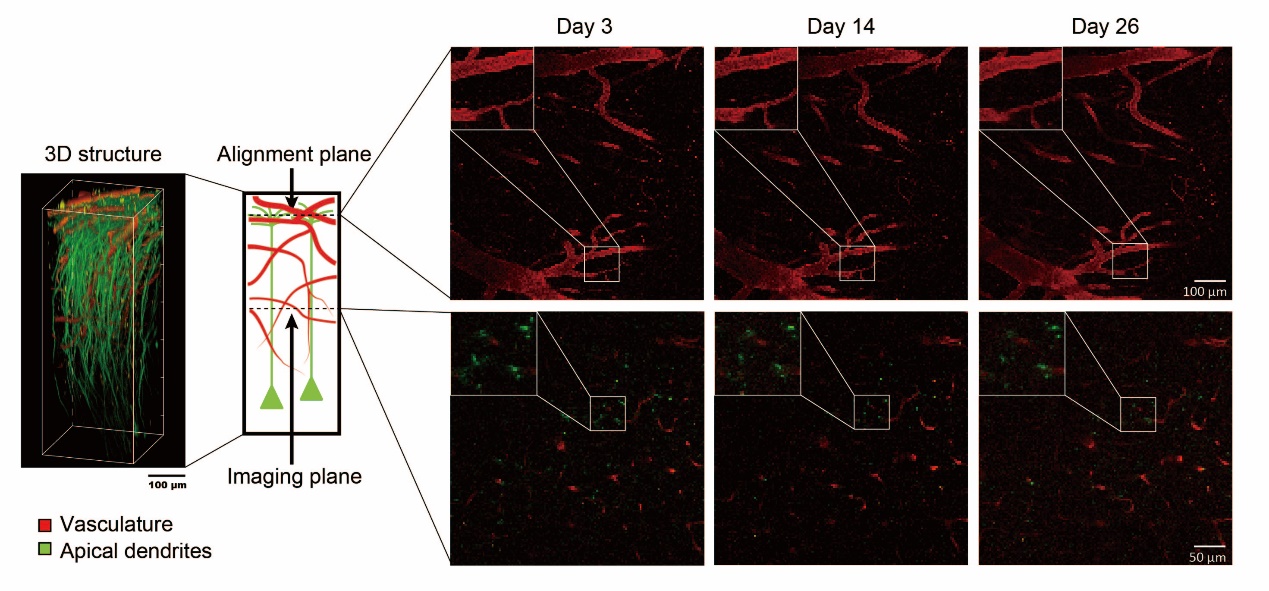

**Figure S1 Imaging and alignment of contralateral-projecting corticospinal neurons (cpCSNs).**

Left: 3D structure of the apical dendrites of cpCSNs and vasculature in the motor cortex. Middle: schematic of alignment plane and imaging plane. Right: example of *in vivo* two-photon images of cpCSNs apical dendrites across days in two planes; images outlined in white, located at the top left, represent magnified versions of the areas highlighted in white within the central region.

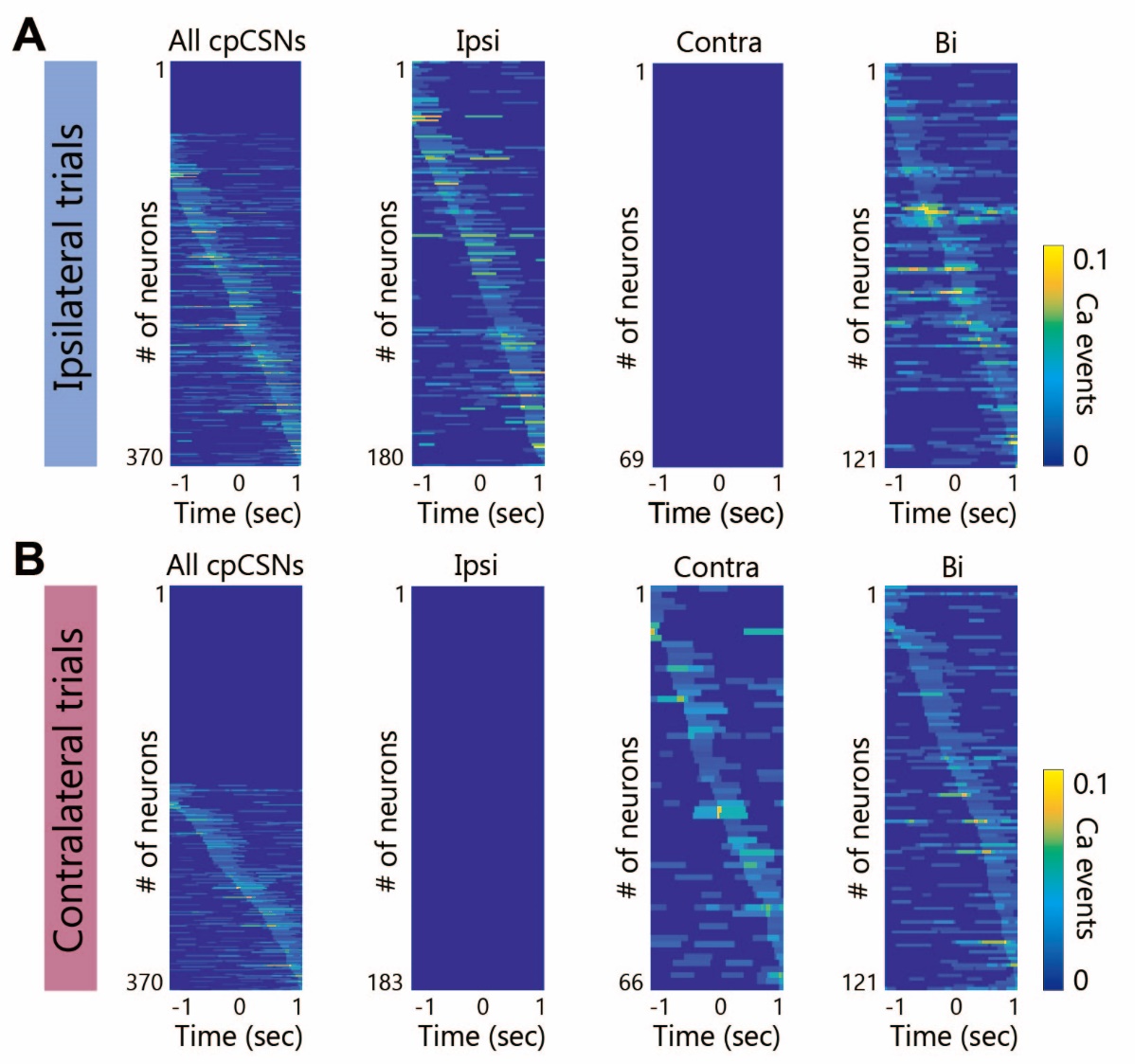

**Figure S2 Examples of activities of contralateral-projecting corticospinal neurons (cpCSNs) during movements.**

Examples of activities of all active, ipsi-preferring, contra-preferring and bi-preferring cpCSNs in ipsilateral trials (top) and contralateral trials (bottom) at different learning stages. Each row represents Ca events averaged across trials of individual cpCSNs, sorted according to their activity level during movements on each side.

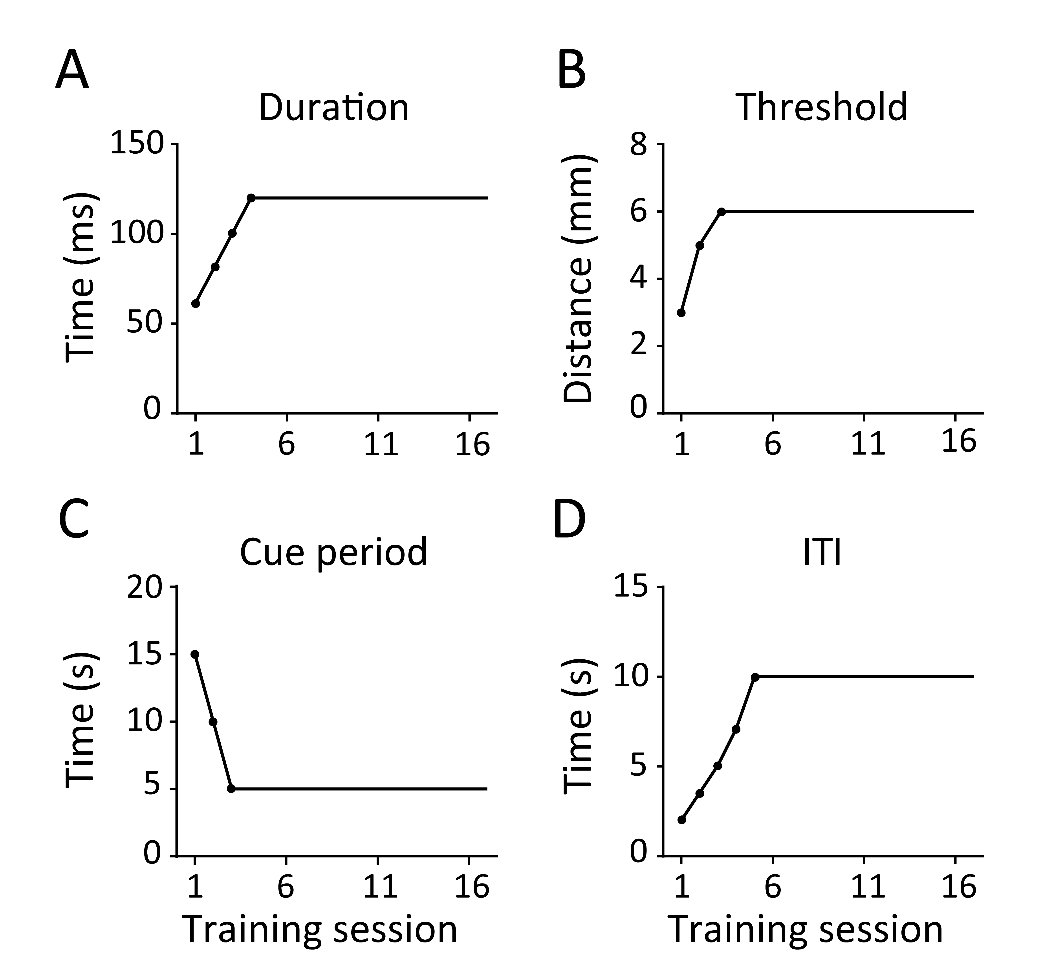

**Figure S3. Behavior task setup in ipsilateral and contralateral sessions during motor learning.**

(A) For both ipsilateral and contralateral movements, the time threshold for lever-press duration was increased during the initial four sessions. (B) For both ipsilateral and contralateral movements, the lever-press distance threshold was raised in the first three sessions. (C) The cue period for lever-pressing was shortened in the first three sessions for both ipsilateral and contralateral movements. (D) The intertrial interval (ITI) was extended for both ipsilateral and contralateral movements over the first five sessions.
